# Structural landscape of engineered multivalent antibody fragments and their application as crystallization scaffolds

**DOI:** 10.1101/2024.10.29.620886

**Authors:** Ashwini Kedari, Liina Hannula, Rupesh Balaji Jayachandran, Ilona Rissanen

## Abstract

Multivalent recombinant antibody fragments, “multibodies”, are produced by fusing antibody VH and VL domains and provide the ability to bind multiple antigens simultaneously. The oligomeric state of a multibody is believed to be determined by the length of the linker region between the V-domains, with longer linkers resulting in diabodies (60 kDa) and shorter linkers leading to the formation of triabodies (90 kDa), tetrabodies (120 kDa), and larger oligomers. In this work, we investigate this design space by engineering multibodies from the sequences of human mAbs CR57 and Imdevimab, and resolve their crystal structures at 2.25 Å and 2.55 Å resolution, respectively. Our results show that despite minimizing the length of the hinge region between the V-domains, these constructs form diabodies. This indicates that linker length is not the sole determinant of a multibody’s oligomeric state, and additional factors such as the mAb origin species and light chain type must also be taken into account when designing multibodies. Moreover, we confirmed that the native paratope of the antibody is well-maintained in the diabody format, and conducted a proof-of-concept trial comparing the crystallization propensity of a diabody versus a Fab in antibody-antigen complex crystallization. Our results show that a diabody can promote crystallization more effectively than a Fab, demonstrating the potential of diabodies as crystallization scaffolds for antibody-antigen complexes.

## Introduction

Antibodies play a fundamental role in molecular recognition within biological systems, crafted by the immune response to selectively bind antigens^1^. The adaptive immune system, triggered by the recognition of foreign antigens, produces antibodies through B cell activation and their differentiation into antibody-secreting plasma cells^2^. Secreted Abs circulate through the bloodstream and tissues, providing protection against pathogens by inhibiting infection and facilitating the clearance of infected cells. Antibodies are also extensively utilized in medical applications and as research tools for biochemistry and molecular and cellular biology. Specifically, monoclonal antibodies (mAbs), which can be recombinantly produced and engineered for diverse applications, offer significant advantages over polyclonal antibodies due to their controlled manufacturing processes, uniform properties, and consistent affinity for specific target antigens^3–5^. mAbs comprise a valued class of biologics applied as therapeutic drugs for treating diseases including cancer, arthritis, and autoimmune diseases^6^. Additionally, mAb fragments are used in structural biology to determine high-resolution structures of antibody-antigen (Ab-Ag) complexes, which often delineate functionally critical sites on the antigens. Engineered mAbs offer the potential to broaden the scope of antibody applications by enhancing affinity^7,8^, specificity^9,10^, and introducing novel functions^11,12^. A detailed understanding of the molecular structure of antibodies is crucial for the successful engineering of improved binding agents.

In their native context, IgG mAbs are Y-shaped proteins (∼150kDa) composed of two identical heavy chains (HCs, gamma (γ) chain in IgG) and two identical light chains (LCs) (Figure 1A)^13,14^. The LC can be of two types, kappa (κ) or lambda (λ). The heterotetrameric structure of IgG is stabilized by extensive protein-protein interactions and disulfide bonds, and both HC and LC chains are further divided into domains that associate to form structurally and functionally distinct regions of the quaternary structure. The canonical mammalian IgG HC is composed of four domains: three constant domains (CH1, CH2, CH3) and one variable domain (VH) (Figure 1A). The LC consists of two domains: one variable (VL) and one constant (CL). The variable domains, as the name suggests, house the hypervariable regions responsible for antigen binding.

**Figure 1.**
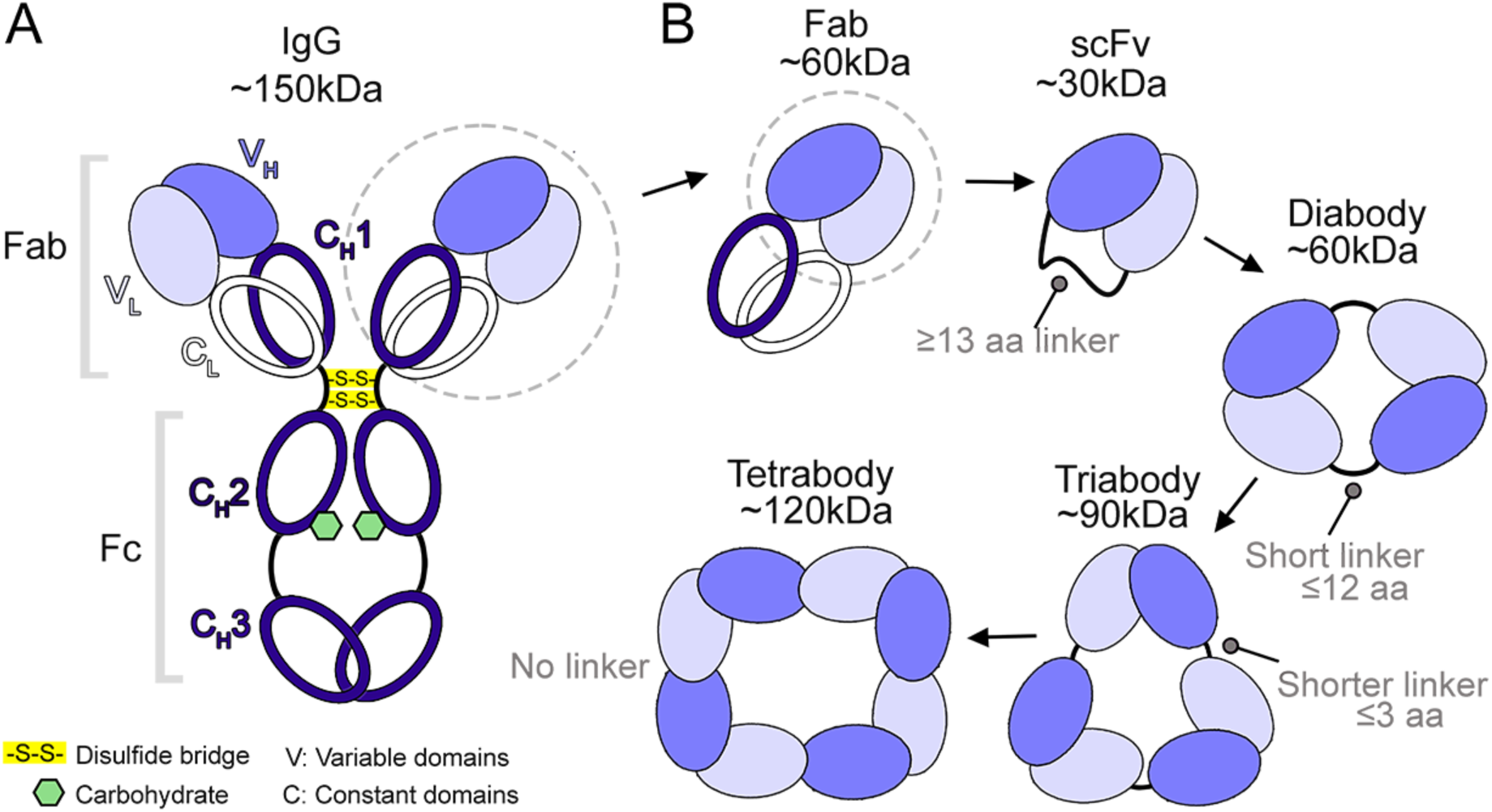
Antibody structure and the principle of engineering multivalent Fv fragments. (A) The canonical mammalian IgG antibody structure (∼150 kDa) with Fab and Fc regions is highlighted. Disulfide bonds (yellow) and the location of the conserved N-linked glycans (green) in the Fc fragment are also shown. (B) Diagram presenting the principles of antibody fragment engineering. In addition to the commonly used Fabs (∼60 kDa), smaller fragments such as scFvs (∼30 kDa), and multivalent forms like diabodies (∼60 kDa), triabodies (∼90 kDa), and tetrabodies (∼120 kDa), can be produced by VH-VL domain fusion, with linker regions of varying lengths between the domains.

Functionally critical regions of the antibody include the Fragment antigen-binding (Fab) regions, which are formed by the pairing of the HC domains VH and CH1 with the LC domains VL and CL (Figure 1). Within the Fab, variable domains VH and VL associate to form the Fv region, which contains the peptides responsible for specific antigen recognition. These consist of six hypervariable loops, known as complementarity-determining regions (CDRs): three in the heavy chain (H1, H2, H3) and three in the light chain (L1, L2, L3)^15,16^. Fabs are connected to another functional component, the fragment crystallizable (Fc) region (formed by CH2 and CH3 of the HC), by largely unstructured hinge regions, which allows the Fabs significant conformational flexibility relative to the Fc^17^. The Fc region can interact with cellular Fc receptors and other immune system components, triggering responses such as phagocytosis and complement activation, and thus provides the effector function profile of the antibody^5,13,18^.

X-ray crystallography is the primary method for achieving atomic-resolution descriptions of Ab-Ag interfaces^19^. However, while full-length mAbs represent the biologically active form of the molecule, the high degree of flexibility between the Fc and Fab regions makes full-length mAbs unsuitable for structure determination by X-ray crystallography. To overcome this limitation and facilitate the crystallization of Ab-Ag complexes, conventional approaches involve truncating mAbs into Fabs or engineering them into single-chain variable fragments (scFvs)^20^. scFvs (∼30 kDa) are produced by fusing the two domains that make up the Fv region, VH and VL, into a single polypeptide chain. In a scFv, the VH and VL domains are connected by a ≥13 amino acid flexible peptide linker (Figure 1) typically consisting of glycine and serine residues^21–23^.

In addition to scFvs, other categories of engineered antibody fragments can also be generated from VH-VL single chain fusions by carefully curating linker length^5,24^. Studies have shown that shortening the linker between the variable domains to 12 amino acid residues or less prevents the variable domains within the same chain from forming the Fv interface, instead inducing the formation of bivalent dimers known as diabodies (∼60 kDa) (Figure 1B)^5,25,26^. Diabodies can also be engineered to be bispecific, enabling them to recognize two different epitopes on the same or different antigens^27–29^. One notable class of bispecific diabodies are bispecific T cell engagers (BiTEs), which simultaneously target CD3 and cancer cell antigens, thereby recruiting T cells to tumor cells and facilitating cytotoxic activity against the targeted tissue^30^. Further reducing the linker size to less than three amino acids, or eliminating it altogether, is reported to lead to the formation of trimers (triabodies, ∼90kDa) and tetramers (tetrabodies, ∼120kDa)^31–34^.

Although strategies for generating multibodies (diabodies, triabodies, and tetrabodies) are available, our understanding of their structural features remains limited, especially for multibodies derived from human mAb sequences. In this study, we address this knowledge gap by determining the oligomeric states and structures of multibodies generated from human mAb sequences using the reported strategies. Furthermore, we postulate that diabodies hold promise as crystallization scaffolds due to their rigidity and the symmetrical arrangement of their two Fv domains, which can facilitate the formation of well-ordered crystals suitable for X-ray crystallography. Here, by comparing the crystallization propensity of diabody−antigen and Fab−antigen complexes, we demonstrate that utilizing a diabody for antigen complex crystallization can offer an effective alternative when traditional methods, such as Fab−antigen complexes, do not produce diffracting crystals.

## Results

### The fusion of human mAb VH–VL domains without a linker results in the formation of diabodies

In this study, we generated multivalent fragments based on the sequences of two human mAbs: CR57, which binds to domain III of the rabies virus glycoprotein (RABV-G)^35^, and Imdevimab (or REGN10987), which targets the receptor binding domain (RBD) of SARS- CoV-2^36^. mAb CR57 was initially isolated from vaccinees and exhibits strong neutralizing activity against multiple RABV strains^35^. Imdevimab was derived from peripheral blood mononuclear cells of human donors previously infected with SARS-CoV-2^36^, and in combination with another human mAb, Casirivimab (REGN10933), it forms the therapeutic cocktail REGEN-CoV (Ronapreve) for the treatment of SARS-CoV-2^37,38^. Here, both CR57 and Imdevimab sequences were utilized to engineer Fab and multibody formats for oligomeric state analysis and structure determination by X-ray crystallography.

For mAb CR57, we first designed a canonical diabody with a five amino acid linker ‘GGGGS’ between the VH and VL domains. In line with reported strategies for diabody formation^24–26^, this design produced a homodimer, as determined by its elution profile in size exclusion chromatography (SEC) (Figure S1). Further, in order to use an engineered mAb CR57 fragment as a crystallization scaffold in our previous study^39^, we generated the same construct without an extraneous linker to increase rigidity, which we expected would result in the formation of tria- and tetrabodies. However, diabody remained predominant, as shown by SEC, which showed a population distribution of 62 % diabody and 36% triabody (Figure 2A). Similarly, the design of the Imdevimab diabody without an extraneous linker between the VH-VL domains produced a predominantly diabody sample (Figure 2B). This propensity for diabody formation was unexpected, given that established strategies for generating multibodies indicate that omitting a linker promotes the formation of larger oligomers^31–34^. To generate a construct with the minimal possible hinge region between the domains and so induce larger multibody formation, we carefully analysed the CR57 diabody structure^39^ (PDB 8R3W; further referred to as CR57 diabody-1). We identified that residues ’SQS’ at the region where the VH domain ends and the VL domain begins to act as a linker in the context of the structure and truncated these residues to create a minimized construct intended to be sterically driven toward forming higher-order oligomers.

**Figure 2.**
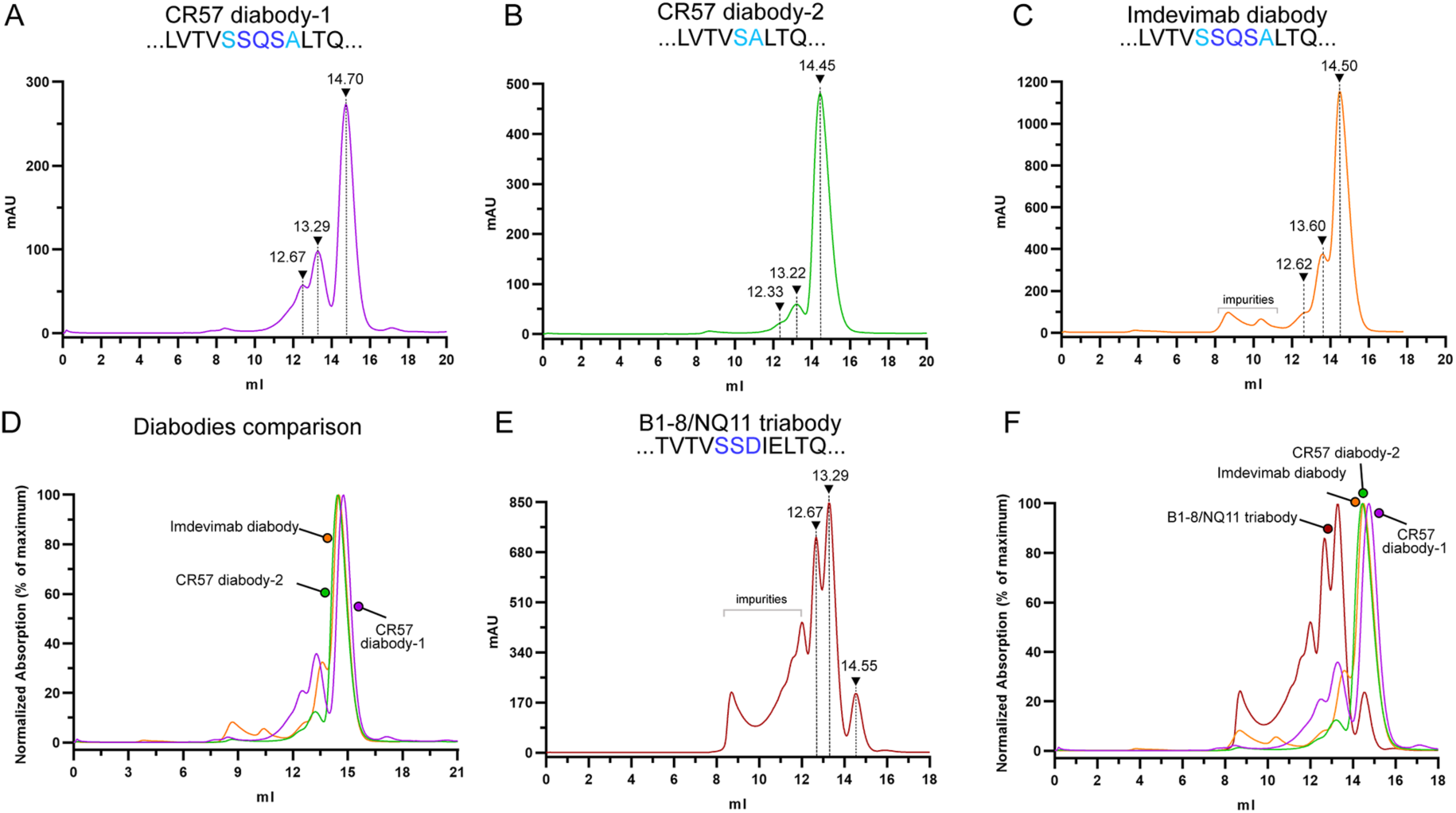
Size exclusion chromatography profiles of the multibodies show distinct oligomeric populations. (A) CR57 diabody-1, (B) Imdevimab diabody, and (C) CR57 diabody-2 all behave as dimers in solution, eluting at 14.7 ml, 14.5 ml, and 14.45 ml, respectively. (D) A comparison of the elution profiles of Imdevimab diabody, CR57 diabody-1, and diabody-2 indicates a similar majority population comprised of diabodies. (E) The B1- 8/NQ11 triabody primarily elutes as a trimer (at 13.29 ml) and larger oligomers, with a minor dimeric population eluting at 14.55 ml. (F) A comparison of all multibody constructs highlights variations in the relative proportion of oligomeric populations. Following the observation that multibody constructs based on human mAbs sequences have the propensity to form diabodies in solution regardless of linker length, we set out to probe the determinants of oligomerization by high-resolution structural characterization. To this end, we resolved the crystal structures of CR57 diabody-2 at 2.25 Å resolution, and Imdevimab diabody at 2.55 Å resolution (Table 1).

**Table 1.** Crystallographic data collection and refinement statistics for the diabodies reported in this study.

|  | CR57 diabody-2 | Imdevimab diabody |
| --- | --- | --- |
| **Data collection** |  |  |
| Space group | $P 6_1 2 2$ | $P 6 2 2$ |
| Cell dimensions |  |  |
| $a, b, c$ (Å) | 69.73, 69.73, 177.58 | 200.55, 200.55, 42.12 |
| $\alpha, \beta, \gamma$ (°) | 90, 90, 120 | 90, 90, 120 |
| Resolution (Å) | 60.38-2.25 (2.29-2.25) | 86.84-2.55 (2.59-2.55) |
| $R_{\text{merge}}$ | 0.112 (1.222) | 0.181 (0.774) |
| $R_{\text{pim}}$ | 0.037 (0.407) | 0.086 (0.379) |
| $I/\sigma I$ | 15.9 (1.6) | 7.5 (1.5) |
| $CC_{1/2}$ | 0.999 (0.845) | 0.980 (0.688) |
| Completeness (%) | 99.6 (100.0) | 99.4 (99.0) |
| Multiplicity | 17.8 (17.9) | 5.3 (5.0) |
| **Refinement** |  |  |
| Resolution (Å) | 60.38-2.25 (2.48-2.25) | 86.84-2.55 (2.71-2.55) |
| No. reflections | 12765 | 16762 |
| $R_{\text{work}} / R_{\text{free}}$ | 0.19 / 0.24 | 0.19/0.22 |
| No. atoms |  |  |
| Protein | 1757 | 1743 |
| Ligand/ion | 24 | 18 |
| Water | 64 | 75 |
| $B$-factors (Å<sup>2</sup>) | | |
| Protein | 53.83 | 42.45 |
| Ligand/ion | 48.35 | 48.85 |
| Water | 48.35 | 47.41 |
| Ramachandran plot (%) |  |  |
| Favored region | 97.80 | 97.38 |
| Allowed region | 2.20 | 2.62 |
| Outliers | 0 | 0 |
| R.m.s deviations |  |  |
| Bond lengths (Å) | 0.002 | 0.008 |
| Bond angles (°) | 0.495 | 0.926 |

\* Values for the highest resolution shell are shown in parentheses

However, SEC revealed that the propensity for diabody formation remained unchanged in the minimized construct, titled CR57 diabody-2 (Figure 2C). Unexpectedly, the minimized construct exhibited a reduced quantity of higher oligomeric populations (89% diabody, 10% triabody) compared to CR57 diabody-1 (Figure 2D).

To date, only one structure of a multibody larger than diabody has been published: that of the B1-8/NQ11 triabody, expressed in *Escherichia coli*^31^. This triabody was generated from two mouse mAbs by fusing the VH domain of mAb B1-8^40,41^ to the VL domain of mAb NQ11 mAb^42^ without any extraneous linker residues between the domains. To determine whether the expression system affects the oligomeric state of this multibody compared to our, CR57 diabody-1, CR57 diabody-2, and Imdevimab diabody constructs, which were produced in human Expi293F cells, we similarly expressed triabody B1-8/NQ11 in Expi293F cells and analysed the oligomeric populations present in the purified sample using SEC. The predominant populations in the triabody B1-8/NQ11 sample consisted of large oligomers (triabodies, tetrabodies, and larger), with only a minor population corresponding to a diabody (Figure 2E). Our SEC results corroborate the findings of the original study^31^, which reported that the B1-8/NQ11 construct produces multiple different oligomeric populations, with the primary population consistent in size with a trimer. This confirms that the differences in oligomerization properties between our constructs and the triabody B1-8/NQ11 (Figure 2F and S2) are not due to the expression system. It also suggests that the conventional strategies for generating larger multibodies by reducing or omitting the linker are not universally applicable to all antibodies, which may be particularly relevant when engineering multibodies from human antibody sequences, as demonstrated in this study.

### Structural analysis reveals that linker length is not the sole determinant of larger order multibody formation

Here, we first present a comparative analysis of structural characteristics of the minimized CR57 diabody-2, the previously reported CR57 diabody-1 (PDB 8R3W)^39^ and triabody B1- 8/NQ11 (PDB 1NQB)^31^.

CR57 diabody-2 crystallized in space group *P* 61 2 2 with one chain present in the asymmetric unit (ASU), potentially indicating a higher degree of rigidity over CR57 diabody-1, which crystallized in space group *P* 21 21 21 with both diabody chains present in the ASU. For our analysis and molecular visualization, the cognate dimer subunit of CR57 diabody-2 was derived from crystal symmetry to complete the biologically relevant diabody assembly. The Fv domain folds of all multibodies (CR57 diabody-1, CR57 diabody-2, and triabody B1- 8/NQ11) are very similar and align well, with root mean square deviation (RMSD) values ranging between 1.0 Å and 1.4 Å for all Fv pairs (Figure S3). The primary difference between diabody and triabody structures is the relative orientation of the Fv domains (Figure 3 and Figure S3). This conformational variability, permitted by flexibility of the regions linking VH and VL domains, is believed to be the primary factor driving higher-order multibody formation in strategies that utilize varying linker lengths to generate diabodies, triabodies, and tetrabodies^31–34^. A detailed examination of these regions reveals that in both CR57 diabody-1 and the B1-8/NQ11 triabody, which share the same design, a segment consisting of three HC and LC residues functions as the linker (Figure 3). By contrast, in the minimized CR57 diabody-2 construct, where these three residues were truncated, the region acting as a linker consists of only two amino acids, 126SA127 (Figure 3B). The longer hinge region peptides of CR57 diabody-1 and the triabody are likely to provide greater rotational freedom compared to the minimized domain fusion of CR57 diabody-2, which is expected to occupy a more limited conformational space.

**Figure 3.**
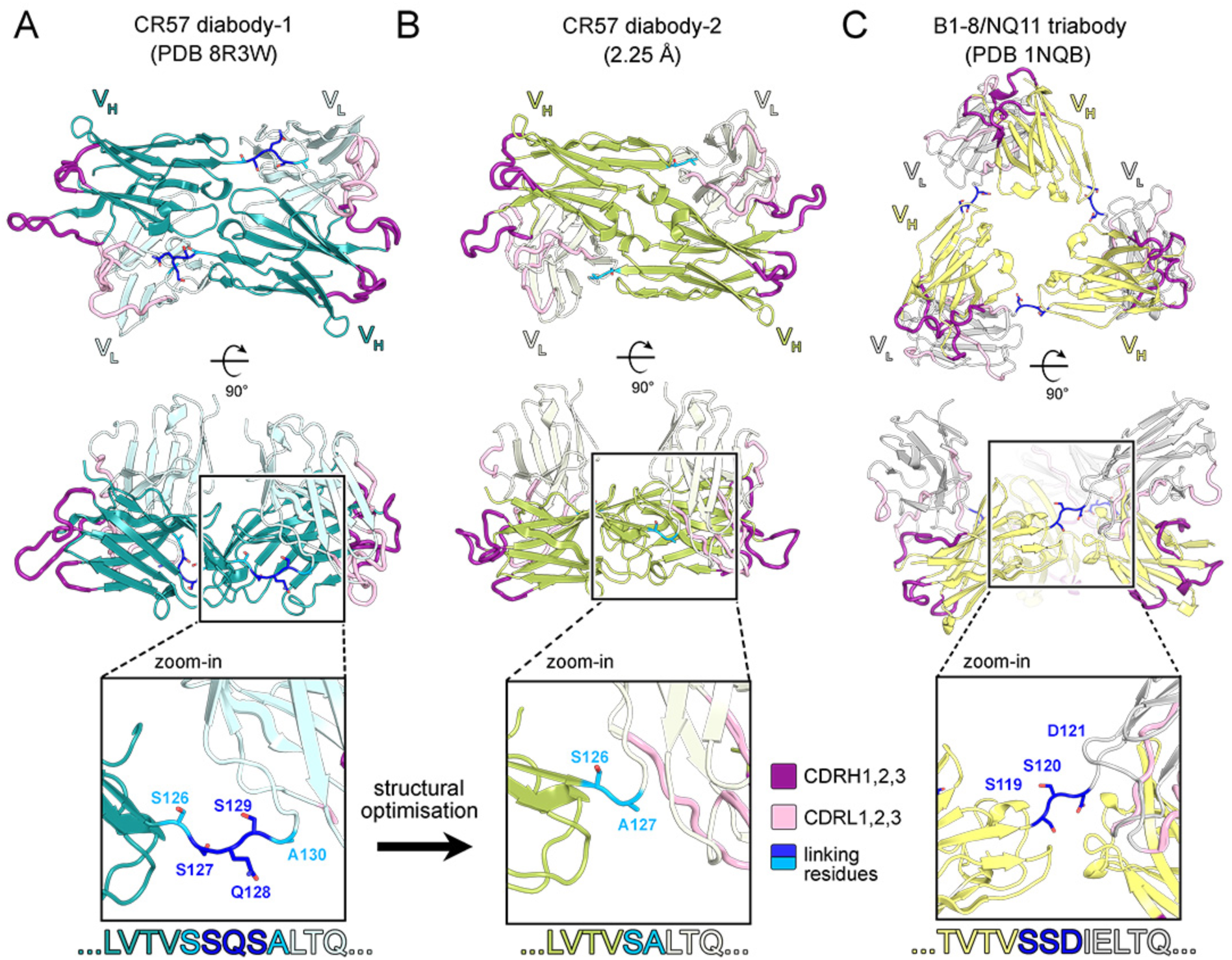
Comparative analysis of the crystal structures of CR57 diabody-1, CR57 diabody-2, and the B1-8/NQ11 triabody. (A) The CR57 diabody-1 crystal structure (PDB 8R3W)^39^, resolved at 2.38 Å, with an inset highlighting the residues “SQS” at the HC and LC domain hinge region that act as a linker in the context of this structure. (B) The optimized CR57 diabody-2 structure, resolved at 2.25 Å, featuring a zoomed-in view of residues “SA”, at the HC and LC domain hinge region. (C) The previously reported B1-8/NQ11 triabody structure (PDB 1NQB)^31^ resolved at 2.0 Å, with a close-up highlighting the residues “SSD”, which form a hinge region of similar length to that observed in CR57 diabody-1.

We note that the diabody nature of CR57 diabody-1 and -2 strongly indicates that multibody formation does not rely solely on linker length and the associated steric constraints. Instead, we postulate that the amino acid residues adjacent to the linkers are likely to play a role, as indicated by the distinct oligomerization profiles of B1-8/NQ11 triabody and CR57 diabody-1, which have the same design but use different antibody template sequences. Specifically, the B1-8/NQ11 triabody was engineered from a mouse antibody with a κ light chain, while CR57 diabodies are based on a human antibody with a λ light chain. Furthermore, as a part of this study, we attempted to generate two diabody constructs based on the human κ light chain antibody Casirivimab (REGN10933)^36^: Casirivimab diabody-1, constructed with no external linkers following the same design as CR57 diabody-1, and Casirivimab diabody-2, following the same minimized design as CR57 diabody-2. No soluble expression of diabodies or larger oligomers was observed for either construct, which further highlights the importance of light chain type and mAb origin species in multibody design. Our results emphasize the need to verify the solution state of engineered multibodies, as similar designs can lead to significantly different outcomes depending on the template sequences used.

### Diabodies promote crystal formation in a proof-of-concept trial with RABV-G domain III and CR57 diabody

While performing crystallization experiments for our previous study^39^ to resolve the structure of the neutralizing mAb CR57 in complex with its targeted viral antigen, RABV-G domain III, we observed that diabody−antigen complexes formed crystals readily. In contrast, no well-diffracting crystals were acquired from domain III complexed with Fab CR57, and the overall tendency of the Fab complex to crystallize seemed lower.

Here, to quantify this difference across a broad range of crystallization conditions, we set up parallel crystallization trials with 672 conditions, using the same antigen complexed with either Fab CR57 or CR57 diabody-1. Crystallization trials were conducted for both complexes at a sample concentration of 5 mg/mL, and since the Fab complex demonstrated greater solubility in pre-crystallization screens, the same array of conditions was also tested with the Fab complex at a concentration of 15 mg/mL. Crystallization screens were set up at room temperature using the sitting drop vapor diffusion method and were automatically imaged. Based on the relative number of conditions that gave rise to visible crystals and precipitation by the 89 days mark, there is a clear indication that the diabody format promotes crystallization more effectively than the Fab (Figure 4). The diabody−antigen complex produced precipitation (217 conditions) and crystal formation, including microcrystals, needles, plates, and 3D crystals (95 conditions) more often than did the Fab−antigen complex (117 and 11 conditions, respectively). Increasing the concentration of the Fab−antigen complex to 15 mg/mL did not increase crystal formation in the same conditions (74 conditions with precipitation, 8 conditions with crystallization). Crystals produced were screened for diffraction at Diamond Light Source synchrotron facility. Only the crystals derived from diabody−antigen complexes exhibited good diffraction properties, leading to the determination of the CR57 diabody−domain III complex structure^39^. These proof-of-concept experiments demonstrate the potential of diabodies as crystallization scaffolds for antibody-antigen complexes and highlight them as a promising alternative to traditional Fab and scFv-based approaches.

**Figure 4.**
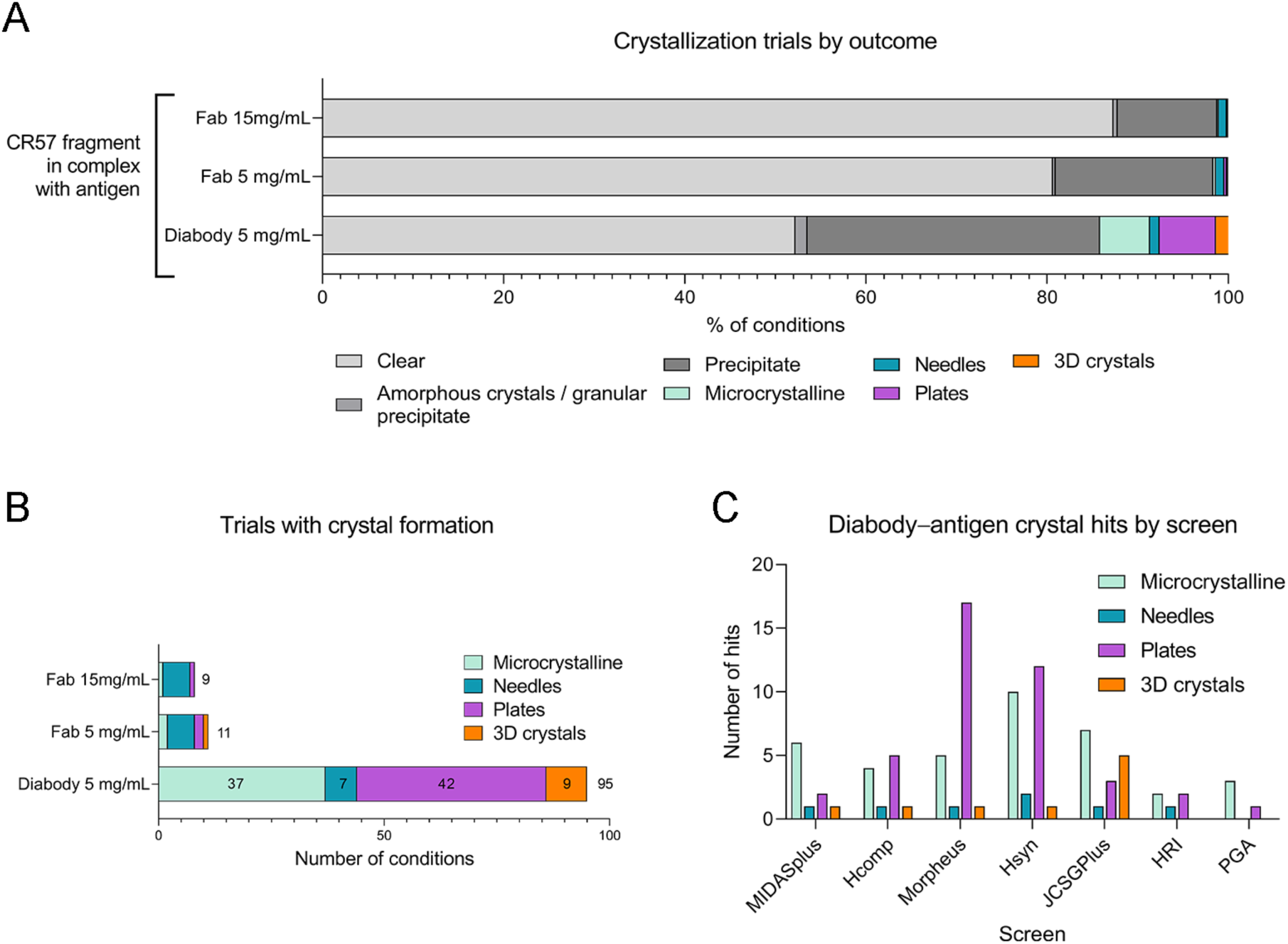
Crystallization of antibody−antigen complexes using Fab and diabody. A) Outcomes from crystallization trials with RABV-G domain III in complex with CR57 Fab at 15 or 5 mg/mL or CR57 Diabody-1 at 5 mg/mL, shown as a percentage of the total 672 trials. B) Number of crystalline hits for each sample. C) Number of diabody−antigen complex crystalline hits in each of the 96-condition crystallization screens tested.

### Structural features of the Fab paratope are preserved in the diabody

Since the CR57 Fab– antigen complex did not yield well-diffracting crystals, a detailed structural comparison between the CR57 diabody–antigen interface and the CR57 Fab–antigen interface could not be performed. To investigate whether a diabody exhibits altered characteristics in the Fv region or at the antigen-binding paratope, we determined the structure of a diabody based on Imdevimab, whose Fab–antigen complex structure was previously resolved using cryo-EM^36^. Imdevimab diabody crystallized readily in a wide range of conditions, forming thick needles that diffracted to 2.55 Å resolution. A single Imdevimab diabody chain was observed in the ASU, and for visualization and analysis of the biologically relevant state of the diabody, the cognate dimer subunit of Imdevimab diabody was derived from crystal symmetry. Superposition of the Imdevimab Fab structure (PDB 6XDG^36^) with Imdevimab diabody shows that the structures align notably well over the Fv domain (Figure 5; RMSD 1.5 Å over 207 aligned residues), which is crucial for maintaining the conformation of the paratope. There were slight differences in the position of the CDR loops, likely due to the absence of bound antigen. Importantly, the original conformation of the antibody paratope is almost exactly maintained in the Imdevimab diabody, even in the absence of the cognate antigen (Figure 5). These findings confirm that the essential features of the antigen-binding region are preserved in the diabody, which is crucial for drawing reliable conclusions about antibody−antigen interface structures derived using a diabody scaffold.

**Figure 5.**
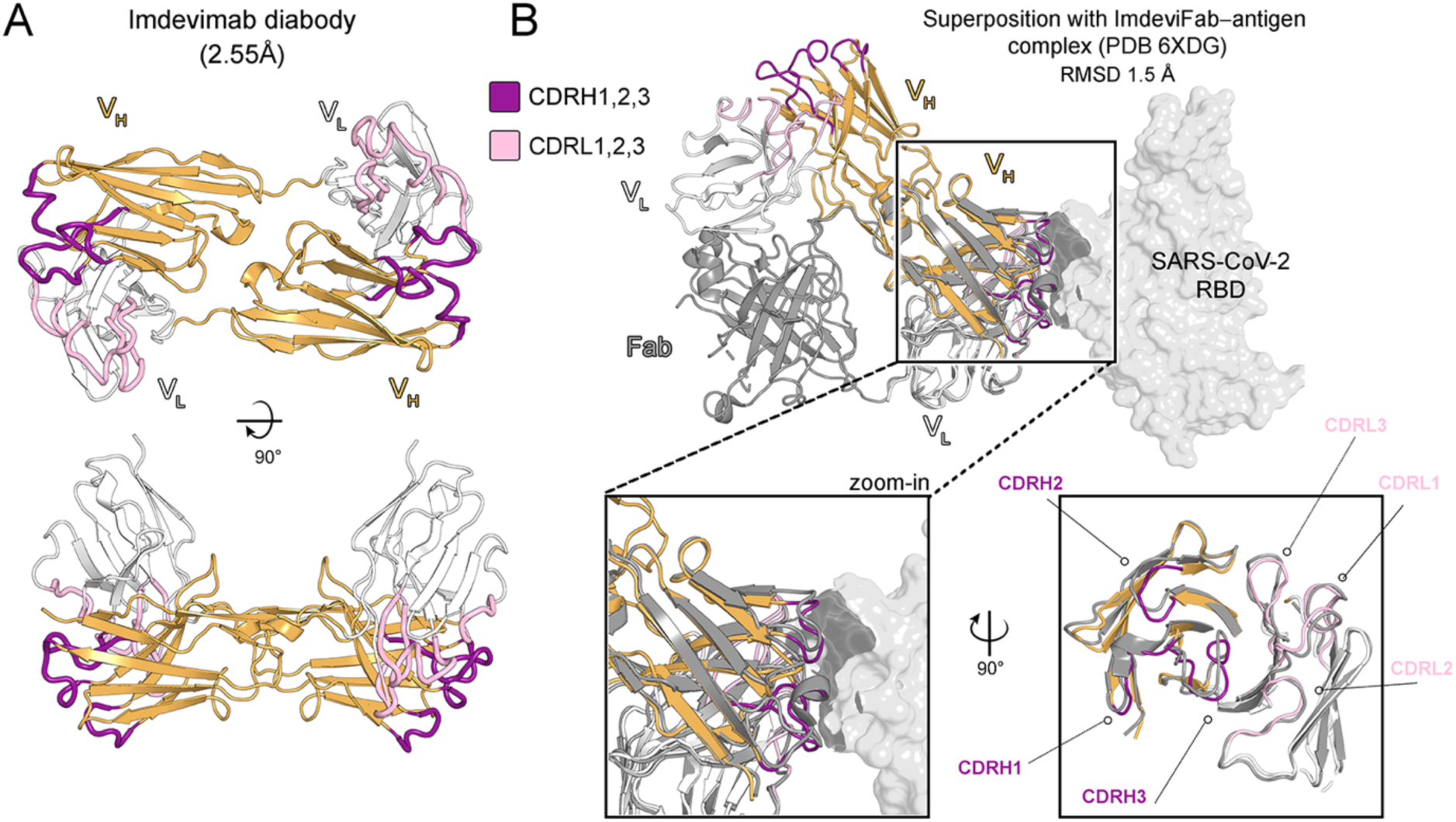
Comparison of Fv and paratope fold between Imdevimab diabody and Fab. A) The structure of Imdevimab diabody was solved to 2.55Å. Diabody model is shown in orange (heavy chain) and white (light chain), with CDR loops highlighted in purple (heavy chain) and pink (light chain). B) Superposition of the Imdevimab diabody model with a previously reported structure (PDB 6XDG^36^) of Imdevimab Fab (grey ribbon) in complex with its antigen, SARS-CoV-2 spike receptor binding domain (grey surface), shows that the overall Fv fold as well as the conformations of the CDR loops are very similar in the diabody and the Fab.

## Discussion

In this study, we explored the structural landscape of multivalent antibody fragments generated from human antibody sequences with no extraneous linker between the VH and VL domains. Two human mAbs, CR57 and Imdevimab, were engineered into multibodies by direct VH and VL domain fusion with the aim of achieving larger-order oligomeric multibodies. Despite the previous success of this design in yielding tria- and tetrabodies^31–34^, our results showed a consistent propensity for these engineered antibody fragments to form diabodies rather than higher-order oligomers. Furthermore, structure-based minimization of the hinge region between the VH and VL domains unexpectedly favoured diabody formation, almost eliminating the larger-order oligomeric population. The preference towards the diabody form in these engineered antibody fragments may stem from the human mAb sequence profile, particularly that of the λ light chain, the impact of which, to our knowledge, has not been previously structurally investigated in multibodies. Based on the observation that multibodies designed using the Casirivimab sequence do not express, we note that the hinge region design supporting the formation of murine κ light chain triabodies and human λ light chain diabodies may be incompatible with the successful folding of multibodies based on human κ light chain sequences. No structures of triabodies or tetrabodies based on human antibody sequences have been reported, underscoring the potential challenges in achieving larger oligomeric multibodies from these template sequences. These findings highlight the need for further research to understand the structural basis of oligomerization in antibody fragments, and to validate the solution states of multibodies generated using established strategies.

X-ray crystallography remains the most prominent method for determining high-resolution structures of Ab-Ag complexes, and crystallization presents a major bottleneck in obtaining structural data. Here, we explored the potential of diabodies as crystallization scaffolds. Earlier reports suggest that symmetric (homo-oligomeric) proteins are more amenable to crystallization than asymmetric (monomeric) proteins^43,44^. Additionally, the bivalent antigen-binding sites of diabodies confer greater avidity compared to monomeric antibody fragments, reducing the likelihood of crystallization occurring without the antigen bound to the engineered antibody fragment. Indeed, our proof-of-concept study, in which we quantified the crystallization properties of Fab–antigen and diabody–antigen complexes, suggests that using diabodies as crystallization scaffolds can enhance the likelihood of obtaining well-ordered crystals compared to Fabs. Furthermore, a structural comparison between an Imdevimab diabody and the previously published Imdevimab Fab structure revealed no conformational changes at the paratope in the diabody format, supporting the use of diabodies as a viable alternative to Fabs for structure determination.

It is important to note that the feasibility of antibody fragment−antigen complex crystallization is likely influenced by the unique features of each complex, including the binding angle of the antibody and the overall size and shape of the antigen, which affect crystal packing. Determining the best antibody format for crystallization could become a part of the optimization pipeline employed in antibody crystallography, and diabodies offer a promising alternative to overcome crystallization issues early on. Overall, our findings contribute to the growing body of knowledge in antibody engineering and pave the way for more effective design and application of antibody fragments in structural biology.

## Acknowledgments

We gratefully acknowledge the facilities and expertise of the HiLIFE Crystallization unit at the University of Helsinki, a member of FINStruct and Biocenter Finland. We thank Diamond Light Source for beamtime (MX37045-1, part of the DLS-CCP4 Data Collection and Structure Solution Workshop 2023, and MX34182-5, part of the Finnish National Protein Crystallography Consortium VI) and the staff of beamlines I03 and I24 at Diamond Light Source for assistance with data collection. Finally, we are thankful to our funders, Research Council of Finland (grant #342988 and #346508 to I.R), University of Helsinki (3-year grant awarded to I.R.), the Finnish Cultural Foundation (grant #00240631 awarded to A.K.), and the Doctoral Programme in Microbiology and Biotechnology, University of Helsinki (1-year salaried doctoral researcher position awarded to L.H.).

## Author Contributions

Conceptualization, I.R.; Investigation, A.K., L.H., and R.J.; Validation, A.K., L.H., and I.R.; Formal Analysis, A.K. L.H., and I.R.; Writing – Original Draft, A.K., L.H., and I.R..; Writing – Review & Editing, A.K., L.H., and I.R.; Visualization, A.K., and L.H.; Supervision – I.R.; Funding Acquisition, I.R.

## Declaration of interests

The authors declare no competing interests.

## Materials and Methods

### Recombinant protein expression and purification

Codon-optimized synthetic cDNA (GeneArt, Life Technologies) encoding the CR57 diabody-1 insert, designed based on the reported sequences of CR57 mAb HC and LC (GenBank AAO17821.1 and AAO17824.1), was cloned into the pHLsec mammalian expression vector (Addgene #99845)^45^ using AgeI-KpnI cloning sites that introduce the vector’s secretion signal and a C-terminal hexahistidine tag into the protein product. The expression construct DNA (50μg) was transfected into the human-derived Expi293F cells (150x10^6^ cells/ml, total volume 50ml) using the reagents and protocols of the Expi293^TM^ expression system kit (Gibco^TM^, ThermoFisher Scientific, catalogue no.14635). The cells were cultivated on an orbital shaker at 37°C with 5% CO2 for six days. The supernatant was collected, clarified by centrifugation and sterile filtered, and the target protein was purified from the supernatant using immobilized nickel affinity chromatography with a 1-ml HisTrap excel column (Cytiva), eluting with 300 mM imidazole. The target protein was further purified by size exclusion chromatography (SEC) using a Superdex 200 10/300 GL Increase column and an ÄKTA go protein purification system (Cytiva) in 10 mM Tris (pH 8.0) and 150 mM NaCl buffer. Templates for the expression of minimized CR57 diabody-2 construct with “SQS” truncation, Imdevimab diabody (design based on sequences reported for GenBank 6XDG_C and 6XDG_A), Casirivimab diabody-1 and Casirivimab diabody-2 (designs based on sequences reported for GenBank 6XDG_B and 6XDG_D), and B1-8/NQ11 triabody (design based on sequence reported for GenBank 1NQB_A) were similarly obtained as codon-optimized synthetic cDNA (GeneArt, Life Technologies), cloned into pHLsec mammalian expression vector (Addgene #99845)^45^, and expressed and purified as described above for CR57 diabody-1.

CR57 Fab heavy- and light chain expression constructs were based on the reported mAb CR57 HC and LC sequences (GenBank AAO17821.1 and AAO17824.1), and the heavy chain construct was truncated at residue 249 to generate a Fab. These sequences were obtained as codon-optimized synthetic cDNA fragments from GeneArt (Thermo Fisher Scientific) and separately cloned into the pHLsec mammalian expression vector (Addgene #99845)^45^. The HC and LC expression vectors were transiently co-transfected (25μg of each plasmid DNA for 50ml of culture volume) into Expi293F cells at a density of 3x10^6 cells/ml using the Expi293 expression system kit (Gibco, ThermoFisher Scientific, catalogue no.14635). Similar protocol for subsequent protein production and purification was followed as described above for the CR57 diabody-1.

RABV-G domain III construct consisting of residues 32-59 and 181-259 (GenBank AAC34683.1), connected by a linker comprised of three glycines, was designed and commercially obtained as codon-optimized synthetic cDNA from GeneArt (Thermo Fisher Scientific). This construct was then cloned into the pURD vector ^46^. A stably expressing protein Expi293F cell line was generated following the method described by Seiradake *et al.* ^47^. Briefly, the domain III/pURD construct and an integrase expression vector pgk-φC31/pCB92 (Addgene #18935) were co-transfected into Expi293F cells which were plated in a 6-well plate (Greiner Bio-One), after overnight cultivation^48^. After another overnight cultivation, the cells were transferred to a T75 flask (Thermo Fisher Scientific). Once confluent, the cells were selected by passaging under 2 µg/ml puromycin for 10 days. After selection, the cells were detached using trypsin, transferred to suspension cell culture in Erlenmeyer shaker flasks (Corning), and scaled-up for protein purification. Protein purification was performed as described previously for CR57 diabody-1. The complex formation of the CR57 diabody-1-domain III was validated by analytical SEC on a Superdex 200 Increase 3.2/300 column (Cytiva) and Aktä pure protein purification system (Cytiva).

### Crystallization and crystallographic data collection

Sitting drop vapor diffusion method at 20°C was used to crystallize samples of CR57 diabody-2 (7 mg/mL) and Imdevimab diabody (10.4 mg/mL). CR57 diabody-2 crystallized in a condition consisting of 0.2M di-ammonium hydrogen citrate and 20 w/v polyethylene glycol 3350, and the Imdevimab diabody crystallized in a condition containing 0.15M malic acid, pH 7, and 20% w/v polyethylene glycol 3500. A cryo-protectant solution (3µl) containing 26% (vol/vol) glycerol was added to the crystals, and the crystals were quickly transferred into liquid nitrogen. X-ray diffraction data were recorded at beamline I03 (λ = 0.97625 Å) for CR57 diabody-2 and at beamline i24 (λ = 0.9999 Å) for Imdevimab diabody, at Diamond Light Source synchrotron, United Kingdom.

For the comparative crystallization trials performed with RABV-G domain III in complex with Fab CR57 (5 mg/mL and 15 mg/mL) or CR57 diabody-1 (5 mg/mL), the following screens were used: Helsinki Complex Screen (https://www.helsinki.fi/assets/drupal/2022-11/hki_complex_screen.pdf), Helsinki Random Screen I (https://www.helsinki.fi/assets/drupal/2022-11/random_i.pdf), Helsinki Synergy Screen (https://www.helsinki.fi/assets/drupal/2024-04/Helsinki_Synergy.pdf), JCSGPlus (MD1-40, Molecular Dynamics), MidasPlus (MD1-107, Molecular Dynamics), Morpheus (MD1-47, Molecular Dynamics), and PGA (MD1-51, Molecular Dynamics). The plates were imaged automatically in the Formulatrix imager at 20 hours, 44 hours, 4 days, 9 days, 14 days, 24 days, 44 days, and 89 days. At the 89 days mark, the droplets were visually inspected, and the crystallization outcomes were scored as one of the following seven categories: clear, precipitate, amorphous crystals or granular precipitate, microcrystalline, needles, plates, and 3D crystals.

### Structure determination

Data were indexed, integrated, and scaled with XIA2 using the dials pipeline^49^. The structure of the CR57 diabody-2 was determined by molecular replacement in Phenix-MR^50^, using a homology model of CR57 Fv generated using the SWISS-MODEL server^51^ based on PDB 5GRX^52^ as the search model. Structure refinement was performed by iterative refinement using Phenix^50^. Coot^53^ was used for manual rebuilding, and MolProbity^54^ was used to validate the model. Structure determination for Imdevimab diabody followed the same process, with the search model for molecular replacement generated from the Imdevimab Fab model (PDB 6XDG^36^) by removing Fab constant regions in Coot, resulting in a scFv model. Molecular graphics images were generated using PyMOL (The PyMOL Molecular Graphics System, Version 2.5.0, Schrödinger, LLC) and UCSF ChimeraX^55,56^.

### Accession Codes

Atomic coordinates and structure factors of the CR57 diabody-2 and Imdevimab diabody have been deposited in the PDB (accession codes AAAA and BBBB respectively).

## Notes

### Competing Interest Statement

The authors have declared no competing interest.

