## Supplementary Information for "Structural landscape of engineered multivalent antibody fragments and their application as crystallization scaffolds"

### Items:

**Figure S1.** SDS-PAGE analysis and analytical size exclusion chromatography (SEC) of diabody CR57

**Figure S2.** Comparison of the size exclusion chromatography profiles of CR57 diabody-1 and -2, Imdevimab diabody, and B1-8/NQ11 triabody

**Figure S3**. Structural comparison between the CR57 diabodies and the previously reported B1-8/NQ11 triabody

**Supplementary data 1.** Amino acid sequences of diabody constructs used in this study

Supplementary References

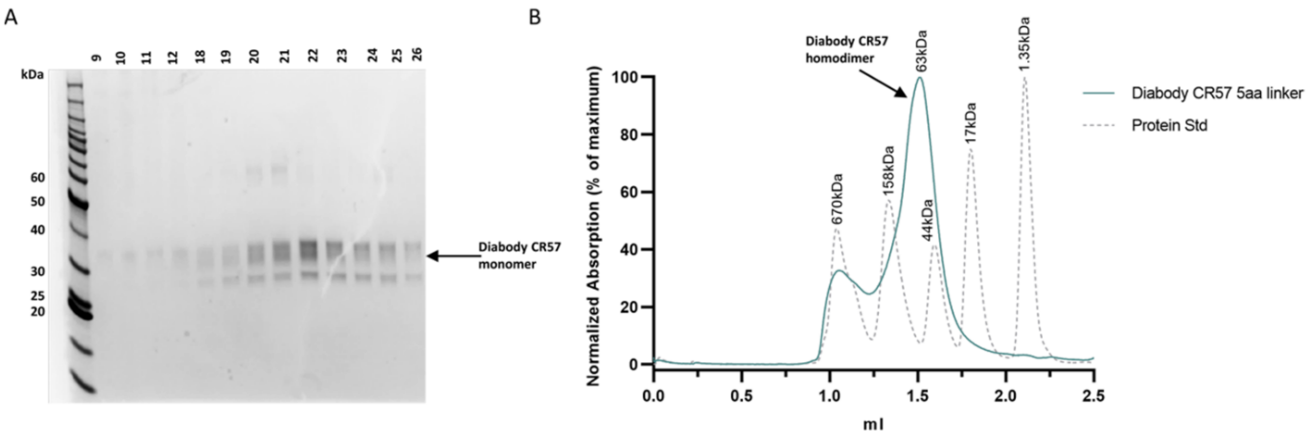


**Figure S1. SDS-PAGE analysis and analytical size exclusion chromatography of the canonical diabody CR57 (related to Figure 2).**

(A) SDS-PAGE analysis reveals a single chain of the CR57 diabody migrating with a molecular mass of ~35 kDa, indicative of a monomer. (B) The size exclusion chromatography results confirm that CR57 exists predominantly as a homodimer in solution, consistent with the expected behavior of a diabody, eluting at a point correlating to a molecular mass of ~60 kDa. The chromatogram of CR57 is overlaid with the elution profile of a protein size standard (Bio-Rad Gel Filtration Standard #1511901). Both the **s**ize exclusion chromatography runs were performed using a Superdex 200 10/300 Increase column and an ÄKTA pure protein purification system (Cytiva) in 10 mM Tris (pH 8.0) and 150 mM NaCl buffer.

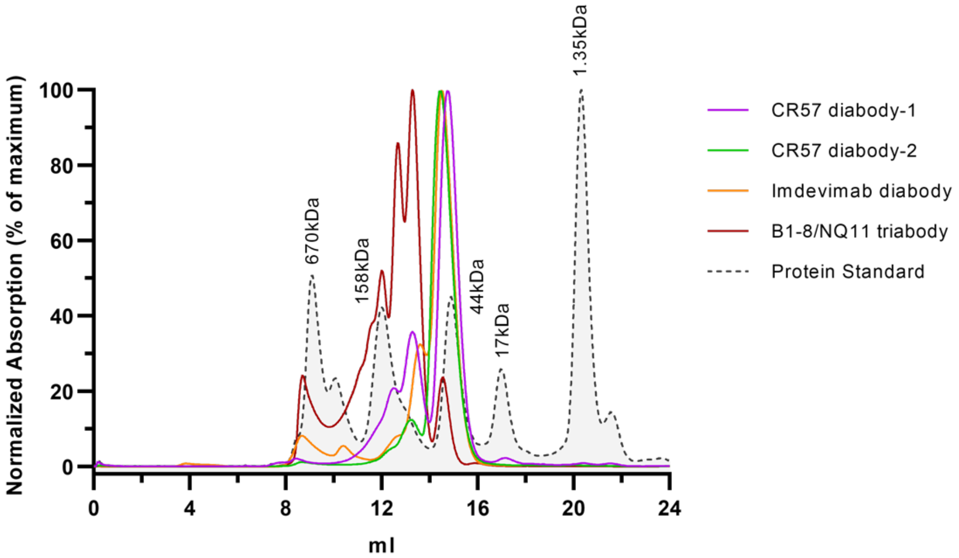


**Figure S2. Comparison of the size exclusion chromatography profiles of CR57 diabody-1 and -2, Imdevimab diabody, and B1-8/NQ11 triabody (related to Figure 2).**

Different multibody constructs are represented by distinct colors: CR57 diabody-1 (purple), CR57 diabody-2 (green), Imdevimab diabody (orange), and B1-8/NQ11 triabody (red). The chromatograms are overlaid with the elution profile of a protein size standard (grey dotted line, with the size of each standard protein indicated; Bio-Rad Gel Filtration Standard #1511901). All size exclusion chromatography runs were carried out using a Superdex 200 10/300 GL Increase column and an ÄKTA go protein purification system (Cytiva) in 10 mM Tris (pH 8.0) and 150 mM NaCl buffer.

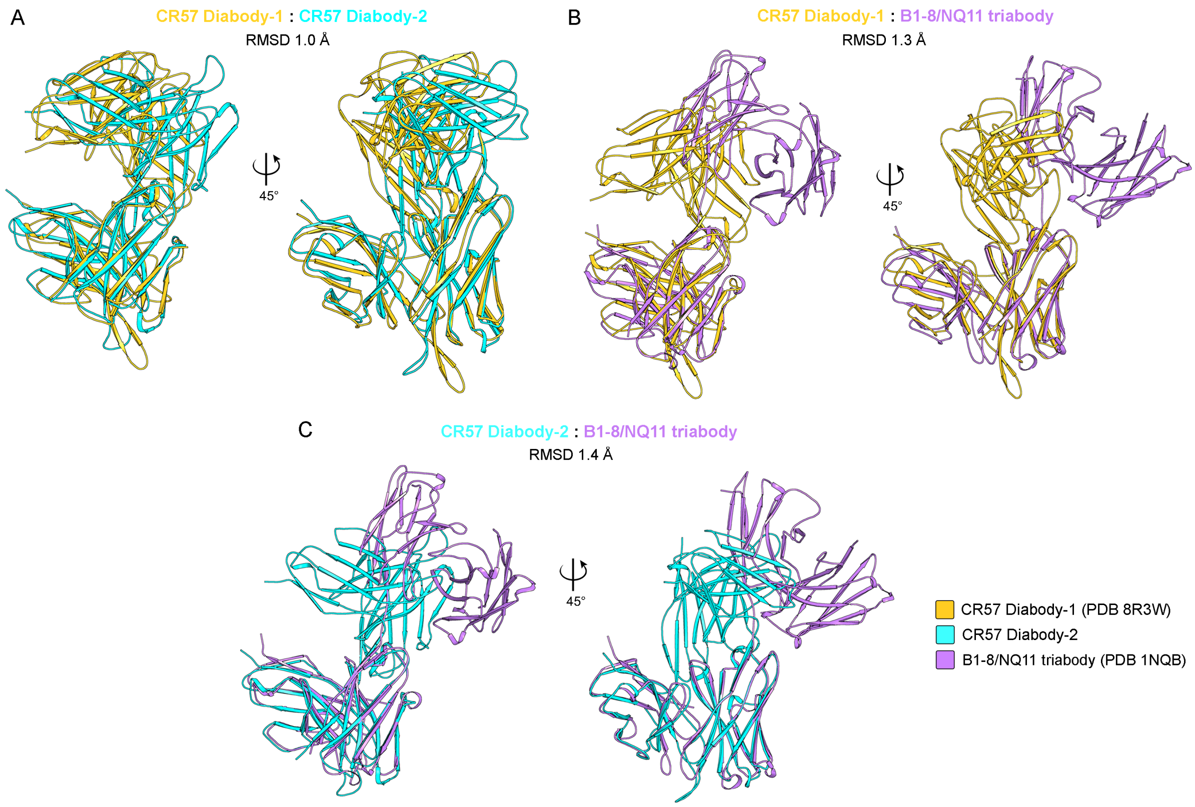


**Figure S3. Structural comparison between the CR57 diabodies and the previously reported B1-8/NQ11 triabody (related to Figure 3).**
(A) Superposition of CR57 diabody-1 (PDB 8R3W)[1], colored gold with CR57 diabody-2, colored cyan. (B) and (C) Superposition of CR57 diabody-1 (PDB 8R3W)[1], colored gold and CR57 diabody-2, colored cyan with the B1-8/NQ11 triabody (PDB1NQB)[2], colored lavender, respectively. All panels show consistent Fv domain folding across the structures, however, the relative orientation of the Fv pairs differs among them.

**Supplementary data 1. Amino acid sequences of the diabody constructs used in this study (related to Materials and Methods).**

**A) CR57 diabody-1**

$V_H$ domain sequence (based on CR57 mAb HC GenBank AAO17821.1) is highlighted in orange and $V_L$ domain sequence (based on CR57 mAb LC GenBank AAO17824.1) is highlighted in grey

**EVQLVQSGAEVKKPGSSVKVSCKASGGTFNRYTVNWVRQAPGQGLEWMGGIIPIFGTAN YAQRFQGRLTITADESTSTAYMELSSLRSDDTAVYFCARENLDNSGTYYYFSGWFDPWG QGTLVTVSSQSALTQPRSVSGSPGQSVTISCTGTSSDIGGYNFVSWYQQHPGKAPKLMIY DATKRPSGVPDRFSGSKSGNTASLTISGLQAEDEADYYCCSYAGDYTPGVVFGGGTKLT VLGQPKAAPSVTL**

**B) CR57 diabody-2**

$V_H$ domain sequence (based on CR57 mAb HC; GenBank AAO17821.1) is highlighted in orange and $V_L$ domain sequence (based on CR57 mAb LC; GenBank AAO17824.1) is highlighted in grey

**EVQLVQSGAEVKKPGSSVKVSCKASGGTFNRYTVNWVRQAPGQGLEWMGGIIPIFGTAN YAQRFQGRLTITADESTSTAYMELSSLRSDDTAVYFCARENLDNSGTYYYFSGWFDPWG QGTLVTVSALTQPRSVSGSPGQSVTISCTGTSSDIGGYNFVSWYQQHPGKAPKLMIYDAT KRPSGVPDRFSGSKSGNTASLTISGLQAEDEADYYCCSYAGDYTPGVVFGGGTKLTVLG QPKAAPSVTL**

**C) Imdevimab diabody**

$V_H$ domain sequence (based on Imdevimab HC; PDB 6XDG[3], chain D) is highlighted in orange and $V_L$ domain sequence (based on Imdevimab LC; PDB 6XDG[3], chain E) is highlighted in grey

**QVQLVESGGGVVQPGRSLRLSCAASGFTFSNYAMYWVRQAPGKGLEWVAVISYDGSNK YYADSVKGRFTISRDNSKNTLYLQMNSLRTEDTAVYYCASGSDYGDYLLVYWGQGTLVT VSSQSALTQPASVSGSPGQSITISCTGTSSDVGGYNYVSWYQQHPGKAPKLMIYDVSKRP SGVSNRFSGSKSGNTASLTISGLQSEDEADYYCNSLTSISTWVFGGGTKLTVLGQPKAAP SVTL**

**D) Casirivimab diabody-1**

$V_H$ domain sequence (based on Casirivimab HC; PDB 6XDG[3], chain C) is highlighted in orange and $V_L$ domain sequence (based on Casirivimab LC; PDB 6XDG[3], chain B) is highlighted in grey

**QVQLVESGGGLVKPGGSLRLSCAASGFTFSDYYMSWIRQAPGKGLEWVSYITYSGSTIYY ADSVKGRFTISRDNAKSSLYLQMNSLRAEDTAVYYCARDRGTTMVPFDYWGQGTLVTVS SDIQMTQSPSSLSASVGDRVTITCQASQDITNYLNWYQQKPGKAPKLLIYAASNLETGVPS RFSGSGSGTDFTFTISGLQPEDIATYYCQQYDNLPLTFGGGTKVEIKR**

**E) Casirivimab diabody-2**

$V_H$ domain sequence (based on Casirivimab HC; PDB 6XDG[3], chain C) is highlighted in orange and $V_L$ domain sequence (based on Casirivimab LC; PDB 6XDG[3], chain B) is highlighted in grey

**QVQLVESGGGLVKPGGSLRLSCAASGFTFSDYYMSWIRQAPGKGLEWVSYITYSGSTIYY
ADSVKGRFTISRDNAKSSLYLQMNSLRAEDTAVYYCARDRGTTMVPFDYWGQGTLVTVS
QMTQSPSSLSASVGDRVTITCQASQDITNYLNWYQQKPGKAPKLLIYAASNLETGVPSRF
SGSGSGTDFTFTISGLQPEDIATYYCQQYDNLPLTFGGGTKVEIKR**
